# Combinatorial binding of semantic information through the sharing of neural oscillatory signals

**DOI:** 10.1101/2023.10.16.562626

**Authors:** Yasuki Noguchi

## Abstract

We comprehend linguistic inputs (e.g. sentence) by retrieving semantic memory of each element (e.g. word) and integrating them. How semantic information is represented and bound as neural (electric) signals is an unsolved issue. I presently used a simple sentence composed of a noun phrase (NP) and a verb (V), comparing human electroencephalography (EEG) responses to a congruent sentence in which the NP and V were semantically related (e.g. “grass grows”) with those to an incongruent sentence (e.g. “a key grows”). In the left temporo-parietal cortex, neural oscillation patterns (8 – 30 Hz) to the second stimulus (V) shifted toward those to the first stimulus (NP), thereby producing coherent (faster and more regular) neural responses to the congruent sentence. No such NP-V interaction was observed in the incongruent sentence. These results indicate that the “semantic unification” as a linguistic concept actually takes place in neural oscillatory signals of the healthy human brain.

## Introduction

In daily conversation, our brain receives a sequence of words as an input, integrating semantic information of those words into the meaning of a whole sentence. This process is called the semantic composition or unification (Hagoort, 2020) and known to play a key role in comprehension of language (Martin and Baggio, 2020; Pylkkanen, 2020). How the brain integrates semantic information across multiple words and phrases is an important but unsolved issue in philosophy, linguistics, psychology and neuroscience (Calmus et al., 2020; Jefferies et al., 2020; Lanzoni et al., 2020; Graessner et al., 2021a).

Electroencephalography (EEG) and magnetoencephalography (MEG) are the methods to non-invasively measure neural activity in the human brain. Their high temporal resolution is suitable to monitor neural responses to linguistic inputs given at natural speed (Peelle and Davis, 2012; Ding et al., 2016; Nelson et al., 2017; Broderick et al., 2018; Gaudet et al., 2020; Heilbron et al., 2022; Kazanina and Tavano, 2023). A well-known EEG response related to semantic composition would be the N400 (Lau et al., 2008), which is evoked by a semantically-incongruent word (critical word) embedded in a sentence. More recent studies investigated neural oscillatory signals related to semantic processing and integration (Weiss and Mueller, 2012; Mollo et al., 2017; Meyer, 2018; Borghesani et al., 2019; Prystauka and Lewis, 2019), although they have reported mixed results; some studies provided data showing a role of gamma rhythm (> 30 Hz) in semantic processing and binding (Bastiaansen and Hagoort, 2015; White et al., 2018; Murphy et al., 2022), while others reported an involvement of alpha (8 – 12 Hz) and beta (13 – 30 Hz) rhythms (Wang et al., 2012; Lam et al., 2016; Teige et al., 2018; Markiewicz et al., 2021; Rempe et al., 2022; Segaert et al., 2022; Zioga et al., 2023).

Most previous studies listed above investigated a change in amplitude (power) of oscillatory signals. In the present study, I analyzed temporal parameters of brain rhythms such as speed and regularity, exploring how the semantic information was integrated between linguistic units as electric signals in the brain. For example, an integration of semantically-related units might induce the successful “merge” of neural activity, producing oscillatory signals of high regularity. In contrast, a presentation of semantically-incongruent word (like a critical word of N400) might evoke the “dissonance” of neural rhythms, resulting in EEG waveforms with low regularity.

Another hallmark of the present study was a use of simple stimuli composed of minimal linguistic units. Although the N400 paradigm typically uses a sentence of around 10 words (Wang et al., 2012; Kielar et al., 2015), such a long sentence induces complex combinatorial operations across those words. Recent neurolinguistic studies therefore used simple sentences with a smaller number of words, focusing on basic combinatorial processes (Bemis and Pylkkanen, 2011; Segaert et al., 2018; Kim and Pylkkanen, 2019; Pylkkanen, 2019; Graessner et al., 2021b; Maran et al., 2022; Hardy et al., 2023). As shown in **Figure 1**, stimuli in the present study were sentences made from a noun phrase (NP) and a verb (V), and participants judged semantic plausibility (congruent/incongruent) of those sentences. For example, “grass grows” is a congruent sentence (CS) in which N (grass) and V (grow) can be semantically integrated, while “a key grows” is an incongruent sentence (IS) violating the selectional restriction. Neural activity for semantic integration would arise selectively to the second stimulus (V) in the CS trials and thus can be detected as an interaction of two-way ANOVA of congruency (CS/IS) × linguistic units (NP/V).

**Figure 1.**
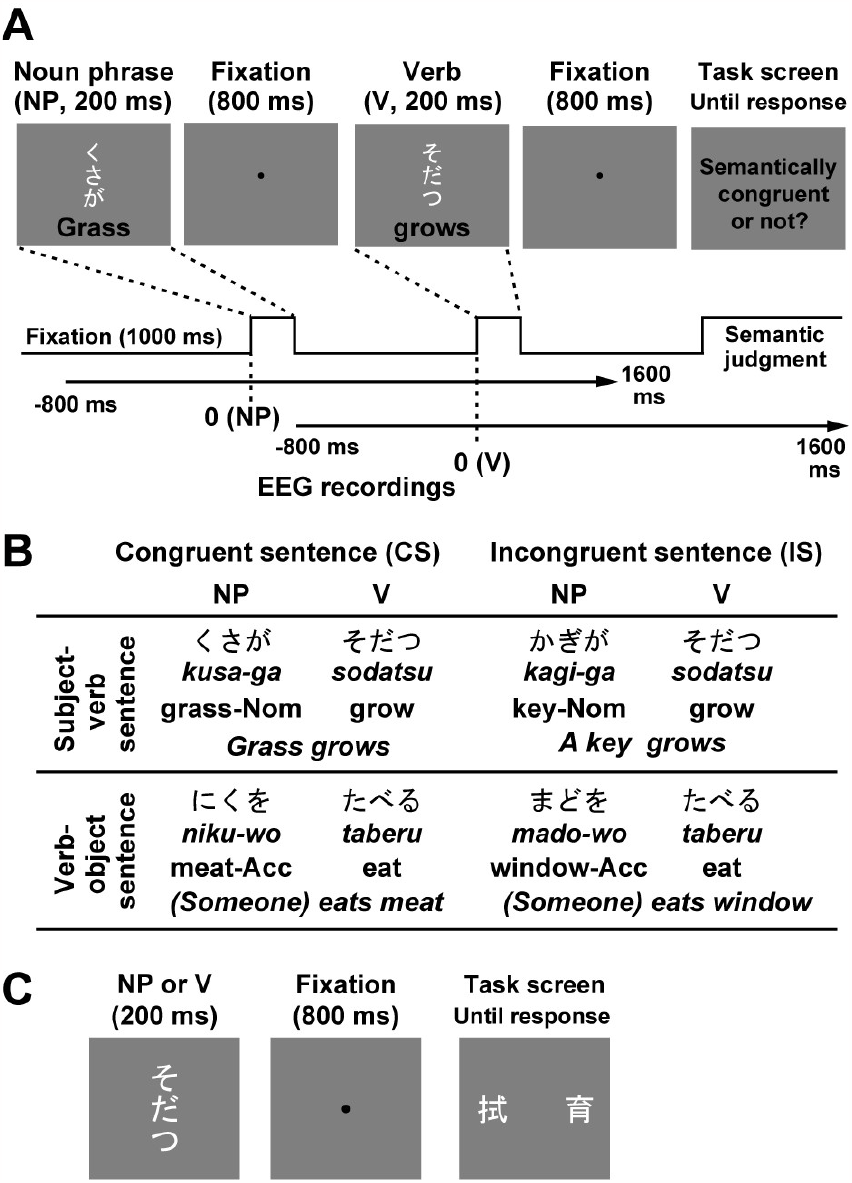
Stimuli and task. (**A**) Semantic task. Each trial consisted of a sequential presentation of a noun phrase (NP, duration: 200 ms) and a verb (V, 200 ms), separated by fixation periods (800 ms). All NPs and Vs were composed of three kana letters (phonograms) in Japanese. Participants judged semantic congruency (interpretability) between the NP and V in the task screen. (**B**) Examples of stimuli. A combination of NP and V formed either a subject-verb sentence (e.g. ‘kusa-ga sodatsu’, meaning ‘grass grows’) or an object-verb sentence (e.g. ‘niku-wo taberu’, meaning ‘(someone) eats meat’), depending on a case particle (‘ga’ or ‘wo’) in the end of NPs. In the congruent sentence (CS), the combination of NP and V was semantically interpretable (e.g. ‘grass grows’ and ‘eats meat’), while it was not in the incongruent sentence (IS, e.g. ‘a key grows’ and ‘eats window’). (**C**) Kanji task. The 120 NPs and 120 Vs used in the semantic task were individually shown. Participants chose one of two kanji letters (ideograms) corresponding to the presented NP or V (phonograms).

## Materials and Methods

### Participants

I recruited 35 healthy participants (native speakers of Japanese, 12 females, age range: 18 - 43), based on results of a power analysis using G*Power 3 (Faul et al., 2007). The type I error rate, statistical power, and an effect size were set at 0.05, 0.80, and 0.5, respectively. Data of 3 participants (1 female) were discarded because of excessive noise in EEG waveforms and replaced by the data of additional 3 participants (1 female). Laterality quotients (LQs) of the Edinburgh Handedness Inventory (Oldfield, 1971) indicated that 33 participants were right-handed (mean: 88.4, range: 26.3 to 100) while two participants were not (-100 and -15.8). All participants had normal or corrected-to-normal visual acuity. I received informed consent from each participant. All experiments were carried out following guidelines approved by the ethics committee of Kobe University, Kobe, Japan.

### Experimental procedures

All visual stimuli were generated and presented using the Matlab Psychophysics Toolbox (Brainard, 1997; Pelli, 1997) and a CRT monitor with a refresh rate of 60 Hz. Each participant performed two tasks in separate experimental sessions. In the first task, he/she viewed a noun phrase (NP) and a verb (V) sequentially presented, judging their semantic congruency as a sentence (semantic task, **Fig. 1A** and **Fig. 1B**). This task has been used in previous studies of fMRI (Suzuki and Sakai, 2003), fNIRS (Noguchi et al., 2002), and TMS (Sakai et al., 2002). Every trial started with a black fixation point (a black circle, diameter: 0.125 deg, duration: 1000 ms) on a gray background. This was followed by a sequence of NP (200 ms) and V (200 ms), separated by a fixation period (800 ms). All NPs and Vs consisted of three kana letters (phonograms) in Japanese. In a half of trials, a noun in the NP was related to V and thus formed a semantically interpretable sentence (congruent sentence or CS. e.g. “grass grows”) while not in the other half (incongruent sentence or IS. e.g. “a key grows”). An order of CSs and ISs within an experimental session was randomized. In the end of each trial (task screen), participants answered semantic interpretability (congruency) of the sentence, pressing one button to CS and another to IS. No time limitation was imposed. Importantly, sentences in all trials were syntactically correct (see “Linguistic stimuli” below), so that participants made their judgment solely based on semantic congruency of NP and V.

In the second task, the 120 NPs and 120 Vs used in the semantic task was individually presented (**Fig. 1C**). In the end of each trial, participants answered a kanji letter (ideogram, imported from China) corresponding to the NP or V, by choosing one of two options on a task screen (kanji task). They pressed one button when the NP/V matched a kanji letter at the left side of the task screen and pressed another when it matched the other kanji at a right side. This conversion from phonogram to ideogram involved the semantic processing but did not need the combinatory processing of NP and V at a sentence level. Each participant performed two sessions of the semantic task (120 trials per session) and two sessions of the kanji task (120 trials per session). An order of those two tasks were counterbalanced across participants.

### Linguistic stimuli

All NPs and Vs had three kana letters. Each NP was made by combining a 2-letter noun and a case particle (“ga” or “wo”) as a third letter. The “ga” is a nominative case particle in Japanese and assigns a role of subject to a noun just before the particle. For example, “kusa-ga sodatsu” is a subject-verb sentence meaning “grass grows” (kusa: grass, sodatsu: grow, upper left in **Fig. 1B**). On the other hand, “wo” is an accusative case particle assigning a role of object. “Niku-wo taberu” is a verb-object sentence meaning “(someone) eats meat” (niku: meat, taberu: eat, lower left in **Fig. 1B**). Note that, while a sentence in English basically requires a subject, Japanese sentence does not. All sentences in the present study therefore had no syntactic error. Combination of 60 nouns with two case particles (“ga”/”wo”) produced 120 NPs. I then chose 120 Vs semantically related to the NPs, making the 120 CS trials. Sentences in IS trials were generated by changing arrangements of those NPs and Vs. All Vs were shown in the present form (not the past form).

It should be noted that the CS and IS trials were balanced except for the semantic congruency. First, NPs and Vs in all sentences were taken from the same list of 240 linguistic units (120 NPs and 120 Vs) so that total visual inputs across all trials were equated between CS and IS. Second, phonological factors were also controlled by fixing a number of letters in each phrase. In Japanese, a number of phonological units (mora) is defined by a number of kana letters in that word. Every NP and V in the present study had 3 moras, which eliminated phonological variations across sentences. Finally, I measured visual similarity between NP and V, comparing those between CS and IS. For each sentence, I counted a number of letters shared by NP and V (common letters). Numbers of those common letters averaged across 120 sentences were 0.108 in CS and 0.1 in IS. No significant difference was observed (*t*(238) = 0.2, *p* = 0.84, Cohen’s *d* = 0.026).

### EEG measurements

Neural activity was monitored with the ActiveTwo system by Biosemi (Amsterdam, Netherlands) from 32 points over the scalp; FP1, FP2, AF3, AF4, F7, F3, Fz, F4, F8, FC5, FC1, FC2, FC6, T7, C3, Cz, C4, T8, CP5, CP1, CP2, CP6, P7, P3, Pz, P4, P8, PO3, PO4, O1, Oz, and O2 (**Fig. 2A**). A sampling rate and an analog low-pass filter were set at 2,048 Hz and 417 Hz, respectively. EEG data were pre-processed by the Brainstorm toolbox (Tadel et al., 2011) for Matlab. First, I removed low-frequency, high-frequency, and fixed-frequency noises from power line using band-pass (0.5 – 200 Hz) and notch (60, 120, and 180 Hz) filters. All waveforms were then referenced with an average potential over the 32 electrodes. EEG responses to NPs and Vs were segmented (epoch range: -800 to 1600 ms relative to a stimulus onset, **Fig. 1A**) and classified into four conditions (CS/IS × NP/V).

**Figure 2.**
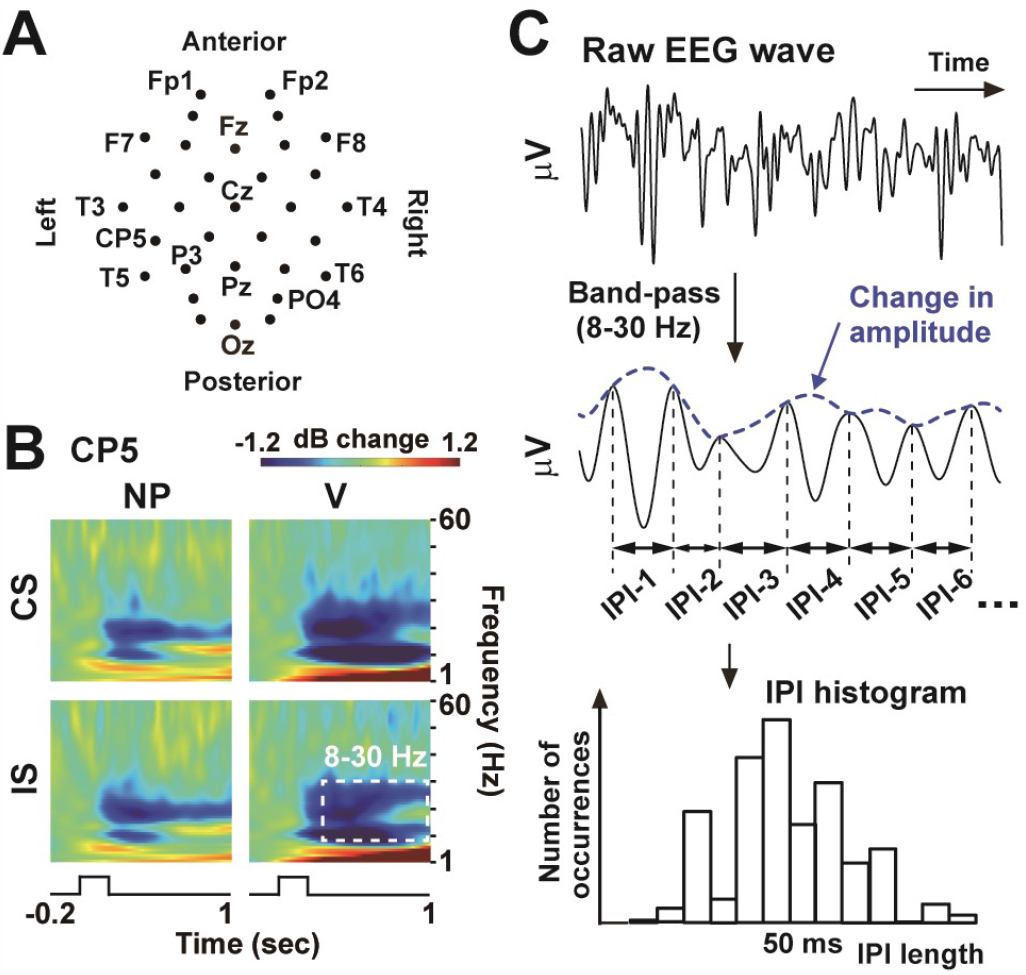
Electroencephalography (EEG). (**A**) Two-dimensional (2D) layout of 32 EEG sensors. (**B**) Time-frequency power spectra (decibel power changes from a pre-stimulus period of -200 to 0 ms, averaged across all participants) at CP5, approximately over the left temporo-parietal junction. Prominent power changes were seen in alpha-to-beta band (8-30 Hz, dotted rectangle). (**C**) Three measures of oscillatory signals; amplitude, speed, and regularity. EEG waveforms at the alpha-to-beta band were first isolated with a band-pass filter. An envelope of the filtered waveform represented changes in amplitude (blue). Speed and regularity were analyzed by measuring time lengths between contiguous peaks of the filtered waveform (inter-peak intervals or IPIs). I pooled all IPIs at 300 – 1000 ms after NP/V onset, depicting a histogram of their occurrences (numbers of IPIs as a function of their lengths). A mean of this IPI distribution provided a measure for the speed of oscillation (a faster rhythm resulted in a shorter mean IPI). The regularity was indexed by a standard deviation (SD) of the IPIs (an oscillation with higher regularity produced IPIs with a smaller SD).

### Analysis of EEG data

I first computed event-related potentials (ERPs) to confirm the N400 response evoked by semantic anomaly. EEG waveforms in each condition were averaged across 120 trials, with the data of a max-min amplitude larger than 150 μV excluded from analyses (target period: -200 to 1000 ms). Numbers of trials that remained after this rejection were 113.6 (CS-NP), 98.3 (CS-V), 113.4 (IS-NP), and 96.8 (IS-V). The N400 would be observed as a differential ERP response to a critical word (V) between CS and IS over the centro-parietal electrodes (Lau et al., 2008; Wang et al., 2012; Panda et al., 2021) such as Cz and Pz.

I then analyzed three measures of oscillatory signals (amplitude, speed, and regularity), especially focusing on the alpha-to-beta band (see **Results** and **Fig. 2B**). First, a band-pass filter of 8 – 30 Hz was applied to EEG waveforms. Change in amplitude of the alpha-to-beta rhythm was measured by identifying an envelope of the filtered waveform with the Hilbert transformation (blue line in **Fig. 2C**). The amplitude waveforms (envelopes) were averaged across all trials in each condition with a baseline set at -200 to 0 ms.

The speed and regularity of oscillatory signals were evaluated with the inter-peak interval (IPI) analysis (Noguchi et al., 2019; Noguchi and Kakigi, 2020). Each peak of the filtered EEG waveform (8 – 30 Hz) was identified, with time lengths between contiguous peaks defined as IPIs (**Fig. 2C**). A mean of IPIs collected over 300 – 1000 ms provided a measure for a speed of neural oscillation, because a slow/fast oscillation resulted in a longer/shorter mean IPI. The analysis period of 300 – 1000 ms was based on the N400 response (see **Results** and **Fig. 3**) showing that the semantic integration of NP and V started from 300 ms after the V onset. The regularity of neural oscillation, on the other hand, was quantified by computing a standard deviation (SD) of the IPIs at 300 – 1000 ms. An oscillation with higher regularity was indexed by a smaller SD, because such a regular waveform produced IPIs highly concentrating around the mean. The mean and SD of IPIs measured in each trial were then averaged across all trials of each condition.

**Figure 3.**
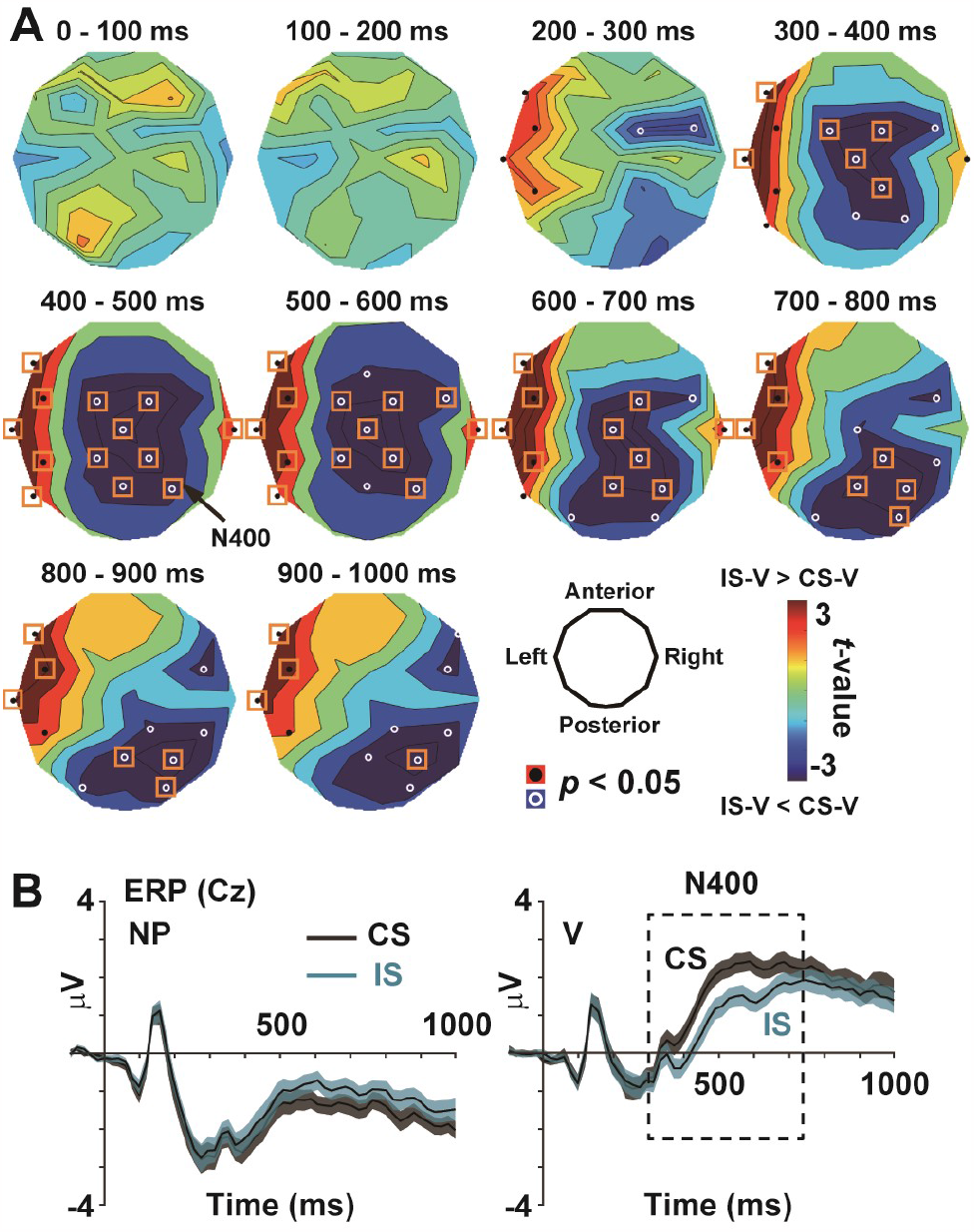
Event-related potentials (ERPs). (**A**) *t*-maps (IS minus CS, following the convention for the N400 studies). Mean ERP amplitudes to V were compared between IS and CS for every 100 ms. Resultant *t*-values (positive: IS-V > CS-V) were color-coded over the 2D layout of 32 sensors. Black and white circles denote sensors showing positive and negative difference of *p* < 0.05 (uncorrected), while orange rectangles denote a significant difference after a correction of multiple comparisons (*p* < 0.05, corrected). Consistent with a previous literature, semantic anomaly in IS elicited the N400 over the centro-parietal region at 300 - 1000 ms (**B**) ERP waveforms at Cz. The left panel shows ERPs to NPs in CS (black) and IS (cyan), while those to V are displayed in a right panel. Background shadings denote standard error (SEs) across participants.

### Statistical procedures

To confirm the N400, I contrasted ERPs to Vs between CS and IS. The ERP waveforms at 0 – 1000 ms were divided into 10 epochs of 100 ms (i.e. 0 – 100 ms, 100 -200 ms, …, and 900 – 1000 ms). For each epoch, an ERP averaged over the 100-ms window (mean ERP) was compared between CS-V and IS-V (N = 35). This comparison was performed for each of 32 sensor positions of EEG, as shown in *t*-maps in **Figure 3A**. A problem of multiple comparisons caused by a repetition of *t*-test for 32 times was resolved by controlling a false discovery rate (FDR) (Noguchi, 2022). A significance threshold was adjusted with the Benjamini-Hochberg correction (Benjamini and Hochberg, 1995) by setting the *q*-value at 0.05. Sensors showing a significant difference after this FDR correction were highlighted by orange rectangles.

The three measures for neural oscillation (amplitude, speed, and regularity) were analyzed in the same way. As the N400, a main contrast was CS-V vs. IS-V (*t*-maps in the middle column in **Fig. 4A-C**). The contrast of CS-NP vs. IS-NP was also made as a control (left column). Finally, since neural responses to semantic integration should be selectively observed in CS-V, an interaction *F*-map of two-way ANOVA (CS/IS × NP/V) was provided in the right column

**Figure 4.**
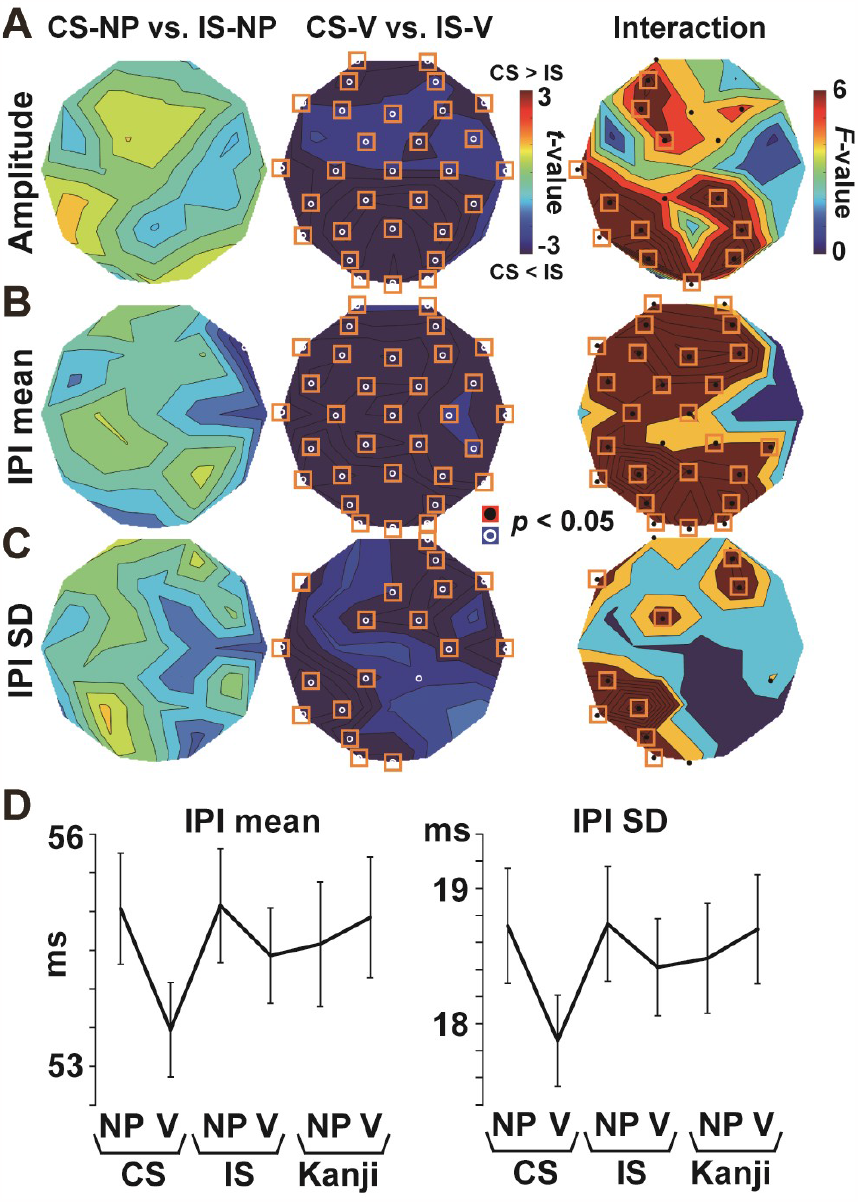
Effect of semantic congruency on oscillatory measures (8 – 30 Hz). (**A**) Amplitude, (**B**) Speed (mean IPI), and (**C**) Regularity (SD of IPIs). The *t*-maps of CS vs. IS (positive: CS > IS) are shown in left (NP) and middle panels (V). Interaction *F*-maps of CS/IS × NP/V are shown in right panels. (**D**) Means and SDs of IPIs averaged across three sensors over the left temporo-parietal cortex (CP5, T5, and P3). Semantic congruency induced smaller (**A**) but faster (**B**) and more regular (**C**) oscillatory signals selective to CS-V.

### Analysis of an IPI distribution

The analysis of oscillatory signals revealed that semantic congruency between NP and V induced an interaction of their neural responses in CS trials (**Fig. 4**). To investigate the details of this interaction, I analyzed histograms of IPIs (occurrences as a function of IPI lengths) and compared them between NP and V (**Fig. 5A**). For each trial, a histogram of alpha-to-beta IPIs was first depicted for NP. An abscissa of the histogram (IPI lengths) was set at 0.49 - 1000 ms, divided into 256 bins from bin 1 (0.49 – 3.91 ms) to bin 256 (996.58 – 1000 ms). Each IPI was assigned to one of those 256 bins depending on its length. In other words, I made an IPI histogram vector for NP (HV_NP_) defined as

**Figure 5.**
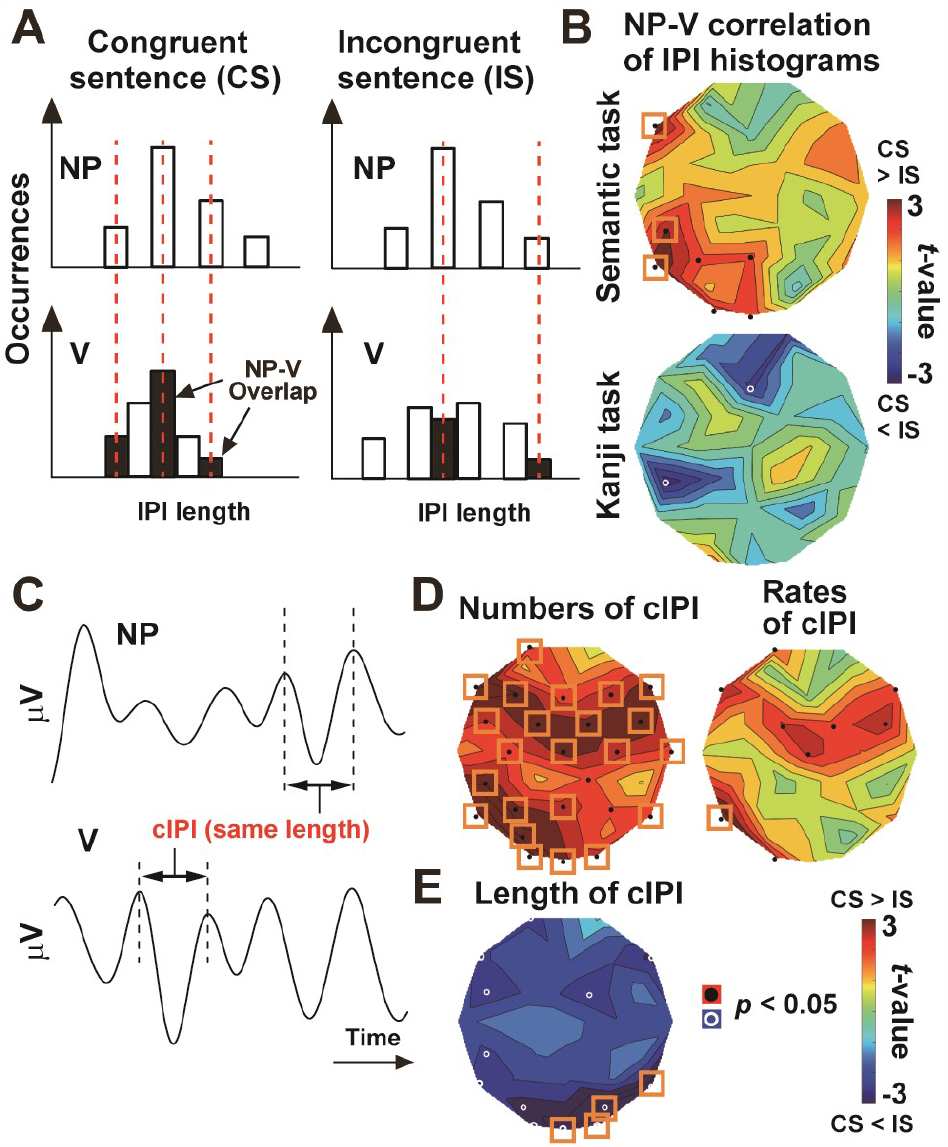
The sharing of IPIs between CS-NP and CS-V. (**A**) Correlation analysis of IPI histograms between NP and V. For each trial, a distribution of IPIs to NP was correlated with that to V. A higher correlation would be observed when the two histograms were more overlapped, sharing the same set of neural oscillations (IPIs). (**B**) *t*-map of NP-V correlation between CS and IS. Sensors over the left temporal cortex showed positive (CS > IS) *t*-values, indicating that IPI histograms of NP and V were more overlapped when they were semantically congruent. No difference was observed when IPI histograms to NPs and Vs in the kanji task (not in the semantic task) were correlated and compared between congruent and incongruent pairs (lower panel). (**C**) Analysis of common IPI (cIPI). The cIPI was defined as the same length of IPI shared by NP and V in the same trial. (**D**) *t*-map for the number of cIPIs (left panel) and their percentage out of all IPIs (right panel) between CS-V vs. IS-V. Semantic congruency produced larger numbers of cIPIs shared by NP and V. (**E**) *t*-map for the lengths of cIPIs between CS-NP vs. IS-NP. The cIPIs in CS were generally shorter than those in IS, indicating that congruent NP and V shared high-frequency oscillatory signals (short IPIs).

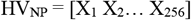

where X_1_ – X_256_ denote numbers (occurrences) of IPIs at bin 1 to 256. Another histogram for V in the same trial (HV_V_) was also made. Finally, I computed a correlation coefficient between the two histograms (HV_NP_ vs. HV_V_). A higher correlation indicates that HV_NP_ and HV_V_ share the same set of neural oscillations (IPIs). This NP-V correlations were averaged across trials and then compared between CS and IS as a *t*-map (**Fig. 5B**, upper panel). Sensors with a significant difference (CS > IS, N = 35) after the FDR correction were shown by orange rectangles.

As a control, I performed the same analysis using data of the kanji task. Based on the same NP-V combinations as the semantic task, the IPI histogram to NP during the kanji task (analysis period: 300 – 1000 ms after the NP onset) was correlated with that to V during the kanji task. Correlation coefficients between HVs for semantically-congruent NP-V pairs were contrasted with those for semantically-incongruent NP-V pairs (**Fig. 5B**, lower panel).

### Representational similarity analysis

The analysis above indicated an involvement of neural oscillatory signals (IPIs) in NP-V integration (see **Results**). On the other hand, a role of IPIs in more basic level (semantic processing of a single word) had remained to be elucidated. I examined this point using the representational similarity analysis (RSA) between linguistic and neural data (Kriegeskorte and Kievit, 2013).

Procedures are summarized in **Figure 6A**. First, I obtained a word vector (1 × 300) for each of 120 Vs (verbs) used in the present study, from the language database online (fastText library https://fasttext.cc/). These word vectors (computed through the word2vec algorithm) were used to make a representational dissimilarity matrix (RDM) reflecting semantic distances among the 120 Vs (linguistic RDM). Each cell in the RDM (dissimilarity index or DI) was computed as 1 - *r*, where *r* was a correlation of word vectors between two Vs. A pair of Vs semantically related to each other (e.g. “grow” and “extend”) produced a low DI (high *r*), while the cell for unrelated Vs (e.g. “grow” and “spill”) was characterized by a high DI. Next, I computed another RDM based on the correlation of IPI histograms (neural RDM). Using the EEG data of kanji task, histograms of IPIs (8 – 30 Hz) for 120 Vs were correlated with each other, producing the second RDM reflecting their neural distances (1 - *r*).

**Figure 6.**
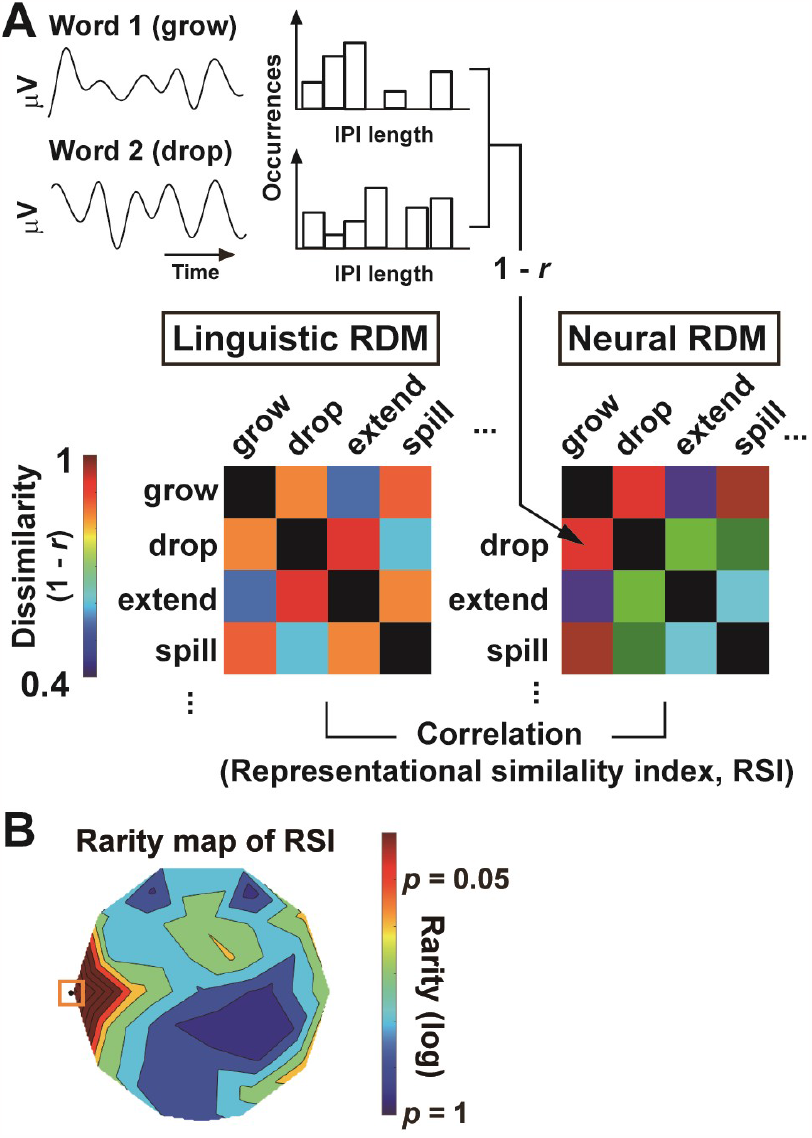
Representational similarity analysis (RSA). (**A**) Procedures. Using word vectors obtained at fastText library, I constructed a representational dissimilarity matrix (RDM) showing semantic distances (1 – *r*) among 120 Vs (verbs) used in the present study (linguistic RDM). Another RDM (neural RDM, 120 × 120) was made based on correlations of IPI histograms and reflected neural distances among the 120 Vs. A high RSI (representational similarity index, a correlation between the two RDMs) indicates that semantically-related words, such as “grow” and “extend”, induced similar distributions of IPIs. (**B**) Results. The left anterior temporal region showed significant (*p* < 0.05 corrected) RSI, indicating that semantic structures (word-word relationships) was represented in IPIs at this sensor (T3). The IPIs were thus related not only to semantic integration of NP and V but also to semantic processing of individual words.

Finally, a correlation between the linguistic (semantic) RDM and neural RDM was computed at each EEG sensor (representational similarity index or RSI). The DIs in bottom-left half of the linguistic RDM were correlated with those of neural RDM, with diagonal DIs (0) excluded from analysis (Chen et al., 2016). A high RSI indicates that semantically-related Vs produced similar distributions of IPIs, showing that semantic information of verbs was represented in neural oscillatory signals. A statistical significance of the RSI was tested through a comparison with random data. By permutating an order of 120 Vs of linguistic RDM for 1000 times, a distribution of RSIs under a hypothesis of null effect was generated. Rarity of RSI in actual data (*p*-value) was estimated as its upper percentile in this random (null) distribution. This rarity map of RSI was shown in **Figure 6B**, with the problem of multiple comparisons resolved by the FDR correction.

## Results

### Behavioral and ERP data

Mean (± SE across participants) accuracy of the semantic and kanji tasks were 95.44 (± 0.39) % and 99.39 (± 0.14) %, respectively. **Figure 3** shows results of ERPs analysis. Difference in mean ERP (IS-V minus CS-V) are shown as *t*-maps over the 32 sensors. A significant N400 response was first observed at 300 - 400 ms, persisting till an onset of the task screen (at 1000 ms). These results ensured that participants detected semantic anomaly embedded in IS.

### Changes in oscillatory signals induced by semantic congruency

**Figure 2B** shows time-frequency power spectra at CP5, approximately located over the left temporo-parietal junction (Sowden et al., 2015; Noguchi and Oizumi, 2018). Both in CS and IS trials, prominent and continuous power changes to NP and V were seen in alpha-to-beta band (8 – 30 Hz). I thus mainly focused on oscillatory signals in this band. Data in other bands (e.g. theta and gamma rhythms) were provided in supplementary materials (**Figure S1**). Results of statistical analyses on oscillatory amplitude are shown in **Figure 4A**. Although a comparison of NPs indicated no difference between CS and IS (left panel), a *t*-map of Vs showed a reduced amplitude of the alpha-to-beta rhythm in CS compared to IS trials (middle panel). A two-way ANOVA of CS/IS × NP/V yielded a significant interaction over the frontal and temporo-parietal cortex especially in the left hemisphere (right panel). Similar results were observed in the mean of IPIs (**Fig. 4B**) and the SD of IPIs (**Fig. 4C**). These results indicate that semantic congruency in CS produced smaller but faster (and more regular) oscillatory responses to V. As shown in **Figure 4D**, this interaction reflected a facilitatory neural processing induced in CS trials, rather than an inhibitory (slow and irregular) response to semantic anomaly in IS trials.

### Attraction of V-response toward NP-response in CS trials

To analyze this neural interaction in details, I correlated histograms of alpha-to-beta IPIs (occurrences as a function of IPI lengths) between NP and V (**Fig. 5A**). If semantic integration in CS can be seen in neural signals, IPIs to NP might be “bound” with those to V, resulting in a substantial overlap of two histograms. Supporting this hypothesis, a correlation coefficient of IPI histograms between NP and V was significantly higher in CS than IS over the left temporo-parietal cortex (**Fig. 5B**, upper panel). Semantic binding was thus characterized by the sharing of the same set of IPIs between NP and V.

As a control, I performed the same analysis using the IPI histograms to NPs and Vs during the kanji task (not the semantic task). No difference in correlation coefficients was observed between semantically-congruent NP-V pairs and semantically-incongruent NP-V pairs (**Fig. 5B**, lower panel). These results suggest that the IPIs to V were attracted toward those to NP only when the information of that NP was stored in working memory (i.e. during the semantic task).

To show this point more directly, I counted number of IPIs shared by NP and V. As shown in **Figure 5C**, a length of each IPI to NP was compared with that to V in the same trial of the semantic task. If a difference in the lengths was within 1.95 ms (within 4 sampling points at 2048 Hz), they were regarded as common IPIs (cIPIs) shared by NP and V. Numbers of cIPIs were found to be larger in CS-V than IS-V (**Fig. 5D**, left), confirming results of the histogram-correlation analysis (**Fig. 5B**). A rate of cIPIs out of toral IPIs (%cIPI) also showed a significant effect of semantic congruency (CS-V > IS-V) over the left temporal cortex (**Fig. 5D**, right). Finally, a comparison of cIPI length (**Fig. 5E**) revealed that cIPIs in the CS trials were generally shorter than those in the IS trials. Semantically-congruent NPs and Vs thus shared IPIs of shorter lengths (high-frequency oscillations) than those in IS, which explained the reduced mean of IPIs selective to CS-V (**Fig. 4B** and **Fig. 4D**).

### A role of IPIs in semantic processing of a single word

Results above showed an involvement of IPIs in semantic integration between linguistic units (NP and V). It remained unclear, however, whether the IPIs also play a central role in semantic processing of a single word. I investigated this point by performing the RSA between linguistic and neural RDMs (see **Materials and Methods**). The linguistic RDM reflected a semantic distance for each pair of 120 Vs used in the present study (**Fig. 6A**). The neural RDM, on the other hand, depicted neural distances among the same set of 120 Vs, which was measured as 1 – *r* where *r* was a correlation of IPI histograms. A high RSI (correlation) between the two RDMs denotes that semantically-related verb pairs produced similar distributions of IPIs, showing the role of IPIs in representing semantic information of individual words.

**Figure 6B** shows a rarity map of RSI. An EEG sensor over the anterior temporal cortex (T3) showed an RSI significantly above a chance (*p* = 0.001). Oscillatory signals (8 - 30 Hz) in this region thus encoded the semantic information of words, showing similar IPIs to semantically-related verbs but dissimilar IPIs to semantically-distant ones.

## Discussion

In the present study, I measured neural activity related to semantic binding. Although most previous studies (Bemis and Pylkkanen, 2011; Flick et al., 2021; Graessner et al., 2021b; Li and Pylkkanen, 2021) focused on the binding process within a phrase (adjective + noun, known as the “red boat” paradigm), I presently investigated the binding between NP and V, as was done in Westerlund et al. 2015 (Westerlund et al., 2015). This was based on a recent finding that sentence stimuli, rather than phrase stimuli, were more suitable to induce changes in neural oscillatory signals (Bai et al., 2022). Results showed that semantic integration in CS trials elicited a reduction in amplitude of the alpha-to-beta rhythm (**Fig. 4A**), consistent with previous studies (Lam et al., 2016; Bonhage et al., 2017; Teige et al., 2018). I further found that semantic binding was characterized by a faster (**Fig. 4B**) and more regular (**Fig. 4C**) neural oscillations over the left temporo-parietal cortex. Finally, despite those prominent changes in oscillatory measures in the CS trials, a correlation of IPI distributions between NP and V was higher in CS than IS trials (**Fig. 5**).

These results suggest that semantic integration induced local changes in IPI distribution of the second stimulus (V) so that it came closer to the first stimulus (NP). As shown in **Figure 5D**, semantic congruency produced larger numbers of cIPIs shared by NP and V. Importantly, this sharing was not observed in congruent NP-V pairs of the kanji task where NPs and Vs were individually presented (lower panel of **Fig. 5B**). This contrast between the semantic and kanji tasks indicates that the sharing of IPIs reflected the NP-V interaction in verbal working memory (WM); the IPIs to V were attracted toward those to NP only when the NP was stored in WM (i.e. the semantic task). In this sense, cIPIs in CS trials might play a role of neural “linker” of WM that persistently retained the sematic information of NPs until it was bound with a subsequent stimulus (V).

As a final analysis, I used EEG data in the kanji task and explored a brain region in which the semantic information of words was represented as the distribution of IPIs. This region was identified in the left anterior temporal cortex (electrode T3, **Fig. 6B**), slightly anterior to the temporo-parietal cortex where the semantic integration took place (T5, CP5, and P3, **Fig. 4A-C**). These results were highly consistent with a previous literature. The anterior temporal lobe (ATL) is known as the semantic “hub” in which various conceptual knowledge was represented as neural signals(Ralph et al., 2017; Farahibozorg et al., 2022). In the present kanji task, participants converted a linguistic unit (NP or V) from phonograms to an ideogram, not needing to integrate the information between units. Indeed, accuracy of the kanji task was very high (99.39 %), much easier than the semantic task (95.44 %). The ATL would play a central role in performing such a simple task. On the other hand, the temporo-parietal cortex, especially a region called the posterior middle temporal gyrus (pMTG), was known as the region for semantic control (Ralph et al., 2017; Jefferies et al., 2020), typically activated by a demanding process like semantic composition. In order to integrate the information of linguistic units (NP and V), the brain had to access semantic knowledge of each unit and connect those under a common concept. The pMTG plays a critical role in such a process involving re-structuring of semantic networks (Ralph et al., 2017). Separate foci of the semantic integration (electrodes T5, CP5, and P3 in **Fig. 4-C**) and the conversion to ideograms (T3 in **Fig. 6**) therefore might reflect a functional division between the anterior and posterior parts of the left temporal cortex.

## Acknowledgments

This work was supported by grants from the Japan Society for the Promotion of Science (KAKENHI: 19H04430) and from The Fukuhara Fund for Applied Psychoeducation Research (Tokyo, Japan) to Y.N. I thank Taeko Kaneda for her technical support. All data supporting the findings of this study are available from Y.N. upon reasonable request. The author declares no competing interests.

## Author contributions

Y.N. developed the study concept, prepared stimuli, collected data, performed data analysis, and wrote the paper.

