## Supplemental Figure S1 for "Combinatorial binding of semantic information through the sharing of neural oscillatory signals"

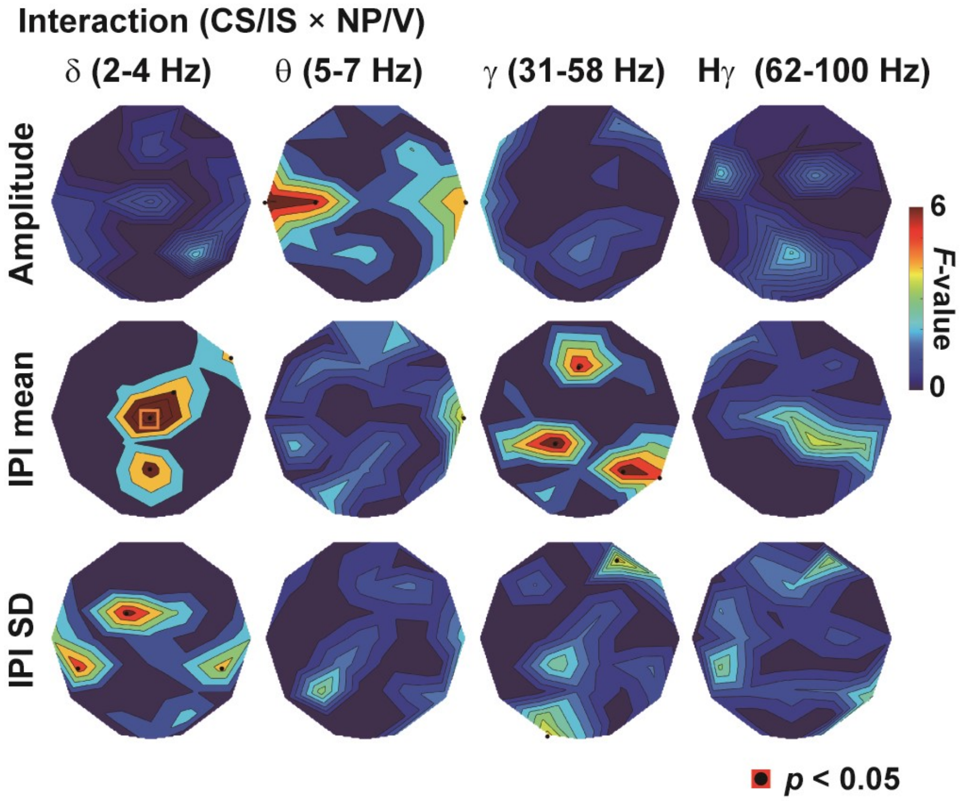


**Figure S1.** Interaction *F*-maps of CS/IS × NP/V in delta (2 – 4 Hz), theta (5 - 7 Hz), gamma (31 – 58 Hz), and high-gamma (62 – 100 Hz) bands. Results of oscillation amplitude, mean of IPIs, and SD of IPIs are shown in upper, middle, and lower panels, respectively. Different from the *F*-maps in alpha-to-beta band (**Fig. 4**), consistent changes across three oscillatory measures were not observed.
